# Depletion of alloreactive B cells by chimeric alloantigen receptor T cells with drug resistance to prevent antibody-mediated rejection in solid organ transplantation

**DOI:** 10.1101/2023.07.25.550322

**Authors:** Anna C. Dragon, Agnes Bonifacius, Murielle Verboom, Michael Hudecek, Constanca Figueiredo, Rainer Blasczyk, Britta Eiz-Vesper

**Author notes:** These authors contributed equally to this work (Rainer Blasczyk, Britta Eiz-Vesper).

## Abstract

In the present study, we developed a novel cell therapy approach to selectively combat antibody-mediated rejection (AMR), a major and unresolved complication after solid organ transplantation (SOT) caused by donor-HLA-specific, alloreactive B cells. Current treatment options including B-cell depletion protocols are inefficient and result in complete loss of humoral immunity. To selectively eliminate alloreactive B cells characterized by corresponding anti-donor-HLA B-cell receptors (BCRs), we engineered T cells with a novel chimeric receptor comprising a truncated HLA molecule fused to intracellular 4-1BB/CD3ξ signaling domains to generate T cells overcoming rejection by antibodies (CORA-Ts). As proof-of-concept, CORA receptors based on HLA-A*02 were shown to bind anti-HLA-A*02 antibodies from the serum of kidney transplant recipients, indicating their suitability to also target the respective membrane-bound anti-HLA-A*02 BCRs on alloreactive B cells. In co-cultures with B-cell lines expressing and releasing anti-HLA-A*02 antibodies, CORA-Ts were specifically activated, released pro-inflammatory cytokines (e.g. IFN-γ, granzyme B), and exhibited strong cytotoxicity resulting in an effective reduction of anti-HLA-A*02 antibody release. A modification of the HLA-A*02 α3-domain within the CORA receptor effectively abrogated T-cell sensitization. Additionally, using CRISPR/Cas9-mediated knockout of a selected binding protein, CORA-Ts were able to resist immunosuppressive treatment to ensure high efficiency in transplant patients. Our results demonstrate that CORA-Ts are able to specifically recognize and eliminate alloreactive B cells, and thus selectively prevent formation of anti-HLA antibodies even under immunosuppressive conditions. This suggests CORA-Ts as potent novel approach to specifically combat AMR and improve long-term graft survival in SOT patients while preserving their overall B-cell immunity.

## Introduction

The only effective treatment of end-stage organ diseases is replacement of the respective organ by SOT. However, one of the major challenges after SOT is AMR, which is considered as outstanding question in transplantation (*1*) and recognized to be even more important than T-cell-mediated rejection (TMR) as most common reason of late allograft failure (*2*). In most cases, donor-specific antibodies (DSAs) directed against mismatched HLA class I and II molecules can be detected in the blood and bind to graft endothelial cells (*3*). Clinically, this is evident by detection of complement deposition, microvascular inflammation with leukocyte infiltration, and endothelial injury, finally leading to irreversible tissue injury and premature allograft failure (*4*). A recent meta-analysis evaluating various follow-up studies of kidney transplantation assessed an incidence of 1-22% for acute AMR and 8-20% for chronic AMR up to 10 years after transplantation (*2*). The incidence is even higher for patients transplanted with other organs or with preformed DSAs. For example, a study evaluating 258 kidney recipients revealed that 30% of patients with detectable DSAs before transplantation developed AMR in a mean follow-up of 5.6 years in comparison with 9% of patients without preformed DSAs (*5*).

Modern immunosuppressive treatment mainly addresses TMR affecting the B-cell alloimmune response only indirectly. Studies showed that such maintenance therapy has only little effects on plasma cells and memory B cells, thereby solely delaying the formation of DSAs (*4*). This led to the development of therapeutic approaches directly targeting B cells and interfering with DSA-mediated destruction cascades, such as depletion of all B cells by the anti-CD20 antibody rituximab. However, clinical benefit of B-cell depletion in chronic AMR could not clearly be proven in randomized controlled trials (*3*). This is hypothesized to be due to the concomitant elimination of regulatory B cell (B_reg_) subpopulations with beneficial effects for allograft tolerance (*6*). Furthermore, since CD20-targeted immunotherapy only has modest effects on plasma cells, proteasome inhibition, for example by bortezomib, has emerged as novel therapeutic approach to induce plasma cell apoptosis. However, in a randomized controlled trial among kidney recipients with AMR, treatment with bortezomib did not prevent the decline of renal function as consequence of progressing AMR (*7*). This was attributed to an only transient decrease in DSA-producing cells followed by an immediate increase in germinal center B cells, again reconstituting memory and plasma B cells (*8*). Further therapeutic approaches including blockade of costimulation and IL-6/IL-6R interference, as well as complement blockade, targeting of CD38 expressed on plasma cells, and apheresis to deplete circulating DSAs have been tested (*3*). However, randomized studies are lacking or not showing significant effects so far, so that there is, until now, no effective treatment in late and chronic AMR. Moreover, unspecific interference with general B-cell functionality in all of these approaches is associated with an increased infection risk in the already immunocompromised transplant recipients, emphasizing the need for a more precise targeting of alloreactive B cells.

T cells engineered with a chimeric antigen receptor (CAR) widely emerged as therapeutic approach to selectively target and eliminate malignant cells in context of different diseases. In hematological malignancies, CAR-T cells exhibited impressive anti-tumor activity prompting their clinical approval in different indications (*9*). Using a similar approach, we aimed to selectively target anti-donor-HLA B cells responsible for AMR via their respective BCRs in the current study to address both, emerging alloreactive naïve as well as pre-existing alloreactive memory B cells in the patient. As a proof-of-concept, we redirected T cells towards alloreactive anti-HLA-A*02 B cells by introducing a novel CAR-like receptor comprising a truncated HLA-A*02 molecule fused to intracellular T-cell signaling domains in order to generate chimeric allo-antigen-specific T <u>c</u>ells <u>o</u>vercoming rejection by <u>a</u>ntibodies (CORA-Ts). Upon recognition of B cells harboring anti-HLA-A*02 BCRs, CORA-Ts were specifically activated and mediated elimination of these B cells, resulting in an efficient reduction of the anti-HLA-A*02 antibody release. Based on these promising results, application of CORA-Ts, after being tested in further pre-clinical models, might serve as innovative approach to specifically combat AMR caused by anti-donor-HLA B cells and effectively improve long-term graft survival in SOT.

## Results

### CORA_sh receptors comprise a properly-assembled HLA-A*02 complex

As alloreactive B cells releasing antibodies with anti-donor-HLA specificity are responsible for AMR and thus the most common reason for late allograft failure (*2*), we developed a cell therapeutic approach to selectively deplete responsible anti-HLA B cells. As proof-of-concept, CORA receptors consisting of a truncated HLA-A*02 α-chain connected to intracellular 4-1BB and CD3ζ-signaling domains were designed to specifically target anti-HLA-A*02 B cells via their anti-HLA-A*02 BCR (fig. 1A). CORA receptor variants differing in their spacer length (CORA_sh with a 12 aa or CORA_lo with a 229 aa spacer domain) were generated to determine the construct with superior HLA assembly and functional abilities. To proof correct expression and folding of the HLA-A*02 molecule within the chimeric receptor, both CORA receptors were transduced into HLA-negative SPI-801 cells and transduced cells enriched via a co-expressed truncated epidermal growth factor receptor (EGFRt) as selection marker (fig. 1B, S1A). By flow cytometry, all CORA_sh^+^ cells could be stained for HLA-A*02 and cell-endogenous β_2_-microglobulin (β_2_m) compared with untransduced SPI-801 cells that did not express both molecules (fig. 1B, S1B,D). On CORA_lo^+^ SPI-801 cells, HLA-A*02 and β_2_m were detectable on 56% and 45% of cells, respectively. As indicated by binding of the conformational anti-HLA class I antibody W6/32, CORA_sh receptors comprised a correctly-folded HLA-A*02-complex loaded with peptide on all cells, whereas CORA_lo receptor were only expressed with the complete HLA complex on 12% of cells, thus indicating that the long spacer impaired complete association of the HLA molecule with β_2_m and peptide by the cell-endogenous machinery (fig. 1B, S1C).

### CORA receptors mediate recognition of patient-derived anti-HLA-A*02 DSAs

To evaluate binding of patient-derived anti-HLA-A*02 antibodies to CORA receptors, sera from seven kidney transplant recipients were selected, from which four had anti-HLA-A*02-DSAs and three had anti-HLA-A*02 antibodies independent from the transplant HLA type. In all seven patient sera, detectable concentrations of anti-HLA-A*02 antibodies could be measured by using Luminex, a multiplexed assay with HLA allele-coupled beads widely used for clinical proof of anti-HLA DSAs (*10*) (fig. 1C). In a crossmatch, these antibodies were examined for their ability to bind to the generated CORA receptors (fig. 1D,E). As expected, anti-HLA-A*02 antibodies present in sera of these patients bound to PBMCs from HLA-A*02-positive healthy individuals and induced complement-dependent cytotoxicity (CDC) as indicated by a significantly increased crossmatch score (*11*) of 7 (dead cells 40-100%). The same sera did not induce CDC towards HLA-A*02-negative PBMCs (score=1; dead cells ≤ 10%) proving HLA-A*02 specificity. Following addition of anti-HLA-A*02 patient sera to SPI-801 cells transduced with CORA_sh receptors, significantly increased CDC (score=6; dead cells 40-80%) was induced, whereas CDC was not increased following addition to untransduced SPI-801 cells. For CORA_lo^+^ SPI-801 cells, anti-HLA-A*02 patient serum slightly increased CDC (score=3; dead cells 10-40%).

Taken together, transduction of CORA receptors comprising a truncated HLA-A*02 α-chain enabled correct assembly of the HLA-peptide complex by the cell-endogenous machinery when connected to a short spacer domain. Such CORA_sh receptors bound to anti-HLA-A*02 DSAs from transplant patients suggesting their suitability to also target the respective membrane-bound BCRs with the same specificity.

**Fig. 1:**
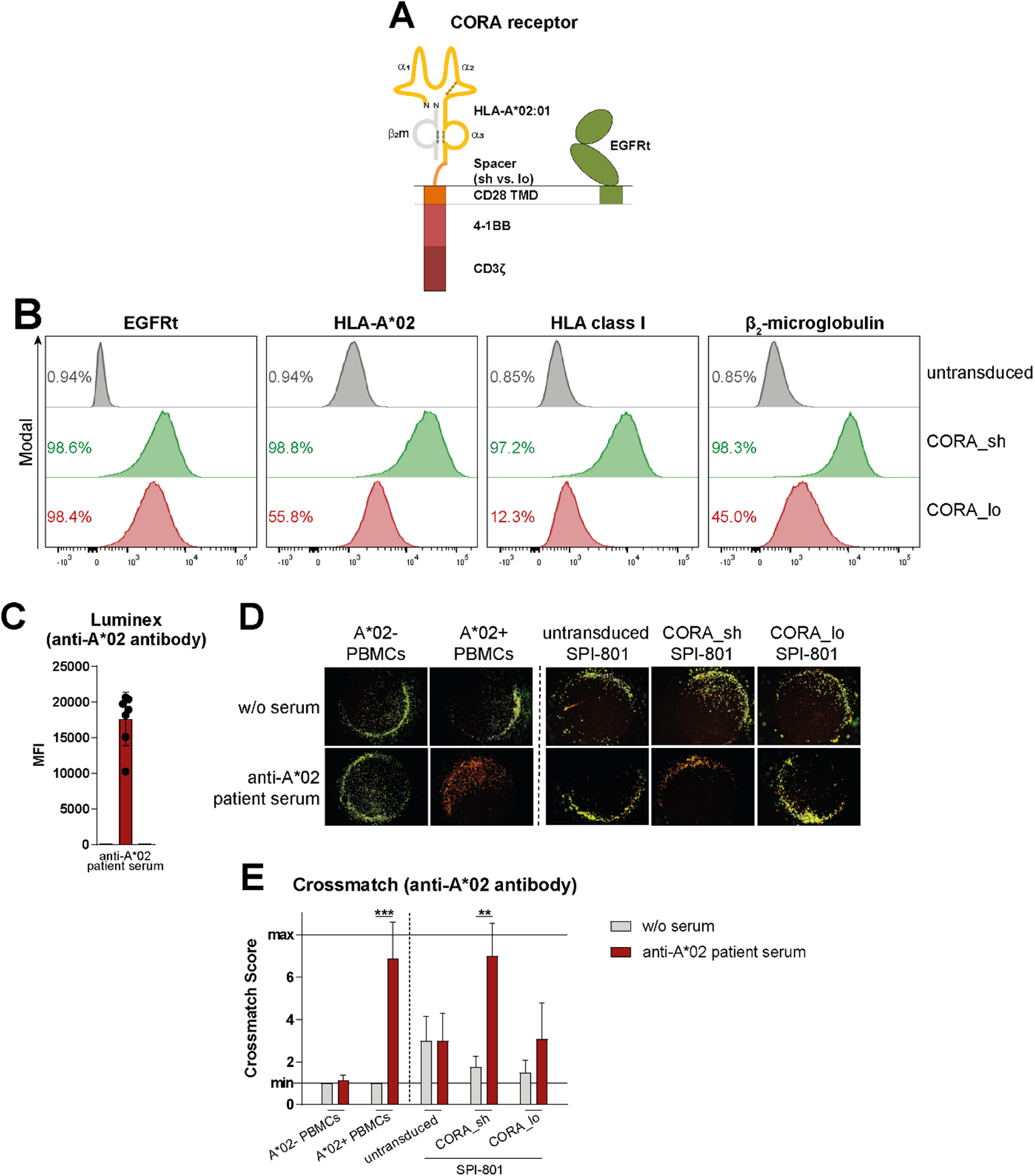
CORA receptors comprising a truncated HLA-A*02 complex mediate recognition of patient-derived anti-HLA-A*02 DSAs. **(A)** CORA receptors comprise HLA-A*02:01 chains α_1_-α_3_, either a short (sh) or a long (lo) spacer domain, a transmembrane domain (TMD) of CD28 and the intracellular signaling domains of 4-1BB and CD3ζ. β_2_-microglobulin (β_2_m, in grey) is not encoded in the vector. A truncated epidermal growth factor receptor (EGFRt) encoded in the same vector serves as transduction and selection marker. **(B)** CORA_sh and CORA_lo receptors were expressed in SPI-801 cells by lentiviral transduction and detected by flow cytometry as shown in representative histograms. Untransduced SPI-801 cells served as control. **(C)** Presence of anti-HLA-A*02 antibody in the serum of kidney transplant recipients was detected by Luminex. Data are shown as scattered dot plot with mean±SD, whereby each symbol represents an individual patient (n=7). **(D, E)** Sera of the same patients were used in crossmatch assays with HLA-A*02-negative or -positive PBMCs from healthy donors, as well as SPI-801 cells transduced with CORA receptors. **(D)** Representative pictures and **(E)** crossmatch scores indicate CDC based on evaluation of viable cells (green) versus dead cells (red) after complement addition. **(E)** Data are shown as mean+SD (n=4-7). Statistical analysis was performed by using Mann-Whitney test. **p≤0.01, ***p≤0.001.

### Hybridoma cells serve as surrogates for anti-HLA B cells releasing anti-HLA DSAs

As targets for evaluation of CORA-Ts, hybridoma cells releasing anti-HLA-A*02 (HB-82), anti-HLA-A/B/C (HB-95) or anti-HLA-B*35 antibodies (TÜ165), respectively, were used and could be shown to express membrane-bound BCRs on 91-98% of cells with comparable mean fluorescent intensities (MFIs) (fig. 2A,B, S2A). Confirming their specificity, HB-82 and HB-95 cells could be stained with eukaryotic HLA-A*02 multimer, whereas TÜ165 cells did not bind to HLA-A*02 (fig. S2B). By Luminex, anti-HLA-A*02 antibodies released by HB-82 cells were detectable in the cell culture supernatant after a short cultivation period and accumulated over time (Fig 2C). The binding specificity of secreted antibodies of all hybridoma cells was confirmed via crossmatch (fig. 2D,E). Similar to anti-HLA-A*02 patient sera, the supernatant from HB-82 cells induced CDC towards CORA_sh^+^ SPI-801 cells in the same extent as towards HLA-A*02-positive PBMCs (crossmatch scores=8; dead cells 80-100%), and crossmatch scores were also slightly increased for CORA_lo^+^ SPI-801 cells (scores=3-4). In line, staining with HB-82 supernatant in flow cytometry revealed antibody binding to 98% of CORA_sh^+^ SPI-801 cells, and to 41% of CORA_lo^+^ SPI-801 cells (Fig 2F, S2C,D). Antibodies present in the supernatant of HB-95 cells revealed the same specificity towards CORA receptors, whereas, as expected, antibodies released by TÜ165 cells did not induce more CDC when compared with respective cells cultured without supernatant (fig. 2D,E). Thus the selected hybridoma cells are suitable surrogates for respective anti-HLA B cells releasing specific anti-HLA DSAs.

**Fig. 2:**
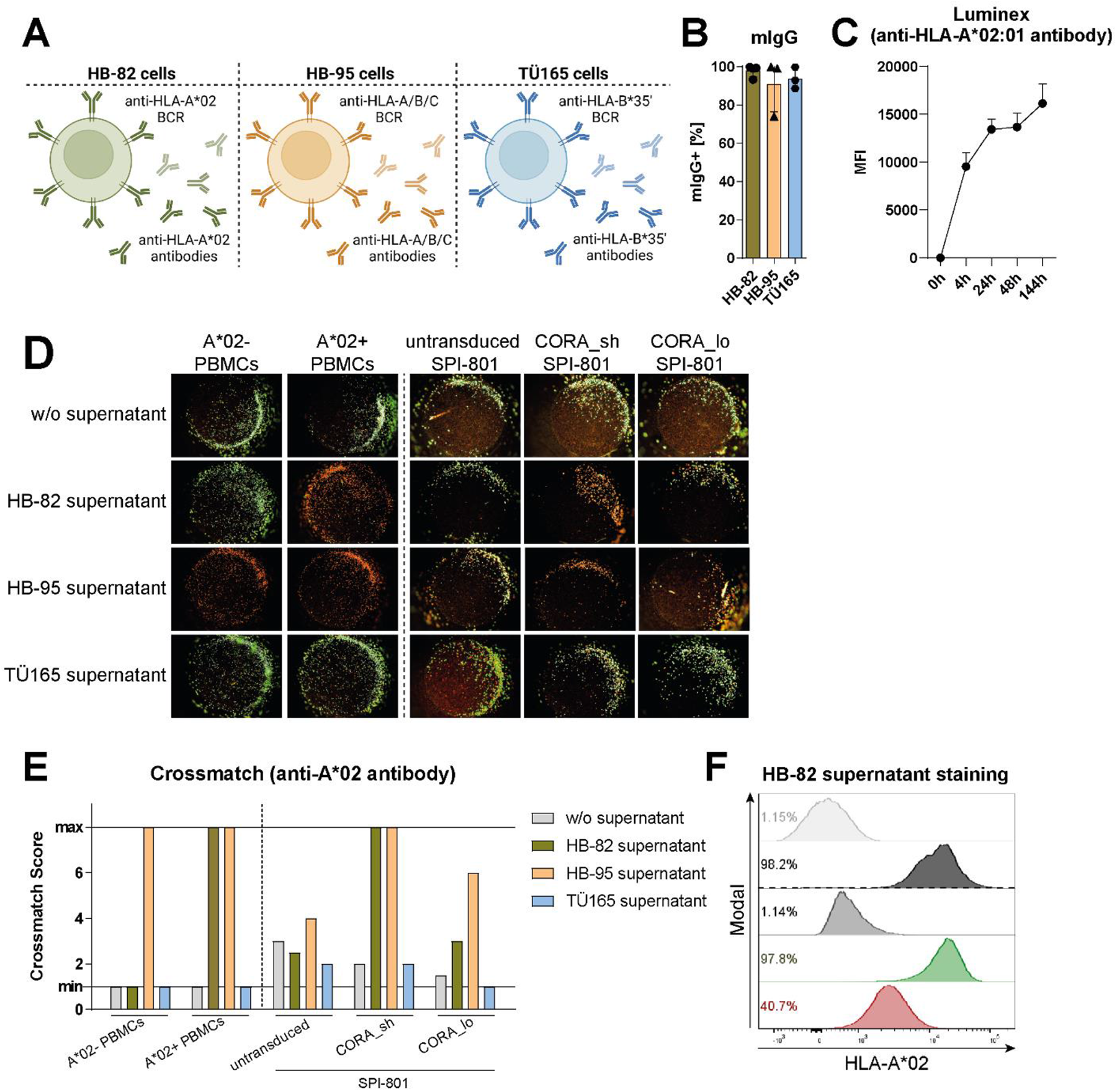
Hybridoma cells serve as surrogates for anti-HLA B cells releasing anti-HLA DSAs. **(A)** HB-82 (anti-HLA-A*02), HB-92 (anti-HLA-A/B/C) and TÜ165 (anti-HLA-B*35 ‘loaded with LPPHDITPY) cells were used as model for anti-HLA-antibody-releasing B cells. Created with BioRender.com. **(B)** Surface expression of murine BCRs was determined by flow cytometry and usage of anti-mouse immunoglobulin G (mIgG) antibody. Data are shown as scattered dot plot with mean±SD, whereby each symbol represents an independent experiment (n=3). **(C)** Release of anti-HLA-A*02 antibody by HB-82 cells was detected by Luminex (n=3-10). **(D, E)** Binding of anti-HLA antibodies present in the supernatant of hybridoma cells to HLA-A*02-negative or -positive PBMCs from healthy donors (all HLA-B*35-negative), as well as to SPI-801 cells transduced with CORA-receptors was assessed by their ability to mediate complement-dependent cytotoxicity (CDC) in crossmatch assays. **(D)** Representative pictures and **(E)** crossmatch scores indicate CDC based on evaluation of viable cells (green) versus dead cells (red) after complement addition. Respective cells incubated without (w/o) supernatant served as viable controls (n=1). **(F)** Cell culture supernatant of HB-82 cells containing anti-HLA-A*02 antibody was used to stain HLA-A*02-negative or -positive PBMCs from healthy donors, as well as CORA_sh^+^ or CORA_lo^+^ SPI-801 cells. MFI: mean fluorescence intensity.

### CORA_sh-Ts mediate effective and target-specific T-cell signaling, activation, cytokine expression and cytotoxicity

To assess the ability of the CORA receptors to recognize anti-HLA-A*02 B cells and initiate T-cell signaling, they were transduced into an established reporter cell line indicating nuclear factor ‘k-light-chain-enhancer’ of activated B cells (NF-κB) and nuclear factor of activated T-cells (NFAT) activation, respectively, by upregulation of reporter fluorophores in flow cytometry (*12*). Following co-culture of either CORA_sh^+^ or CORA_lo^+^ reporter cells with the anti-HLA-A*02 HB-82 cells, activity of both transcription factors was significantly upregulated, whereby the CORA_sh receptor mediated a more pronounced activation (fig. 3A,B). Both receptors were then transduced into primary CD8^+^ T cells isolated from healthy individuals, whereby presence of correctly-assembled HLA-A*02 after enrichment via co-expressed EGFRt was confirmed on 77% of CORA_sh-Ts and 36% of CORA_lo-Ts, respectively (fig. S3A,B). Following co-culture with HB-82 cells, the expression of CD25, CD69 and CD137 as markers for T-cell activation was significantly increased on CORA_sh-and CORA_lo-Ts in a concentration-dependent manner when compared with respective T cells cultured without target cells (fig. 3C-E). In that, the CORA_sh receptor mediated significantly higher T-cell activation than the CORA_lo receptor. In addition, expression of the effector molecules tumor necrosis factor (TNF-)α and granzyme B in CORA-Ts was increased in the same target-specific and concentration-dependent manner (fig. 3F,G). Importantly both, CORA_sh-and CORA_lo-Ts mediated significant cytotoxicity towards HB-82 cells when compared with untransduced T cells as indicated by an elevated release of lactate dehydrogenase (LDH) into the supernatant, whereby CORA_sh-Ts eliminated HB-82 cells significantly more efficiently (fig. 3H).

Thus, CORA receptors with a truncated HLA-A*02 molecule as recognition domain mediate significant effector functionality of engineered T cells towards target cells expressing anti-HLA-A*02 BCRs. In all functionality tests, CORA_sh-Ts showed significantly improved effector functions when compared with CORA_lo-Ts, which is in line with their improved expression of correctly-assembled HLA-A*02 complex as part of the CORA receptor. Hence, the subsequent experiments were focused on CORA-Ts harboring a short spacer domain.

**Fig. 3:**
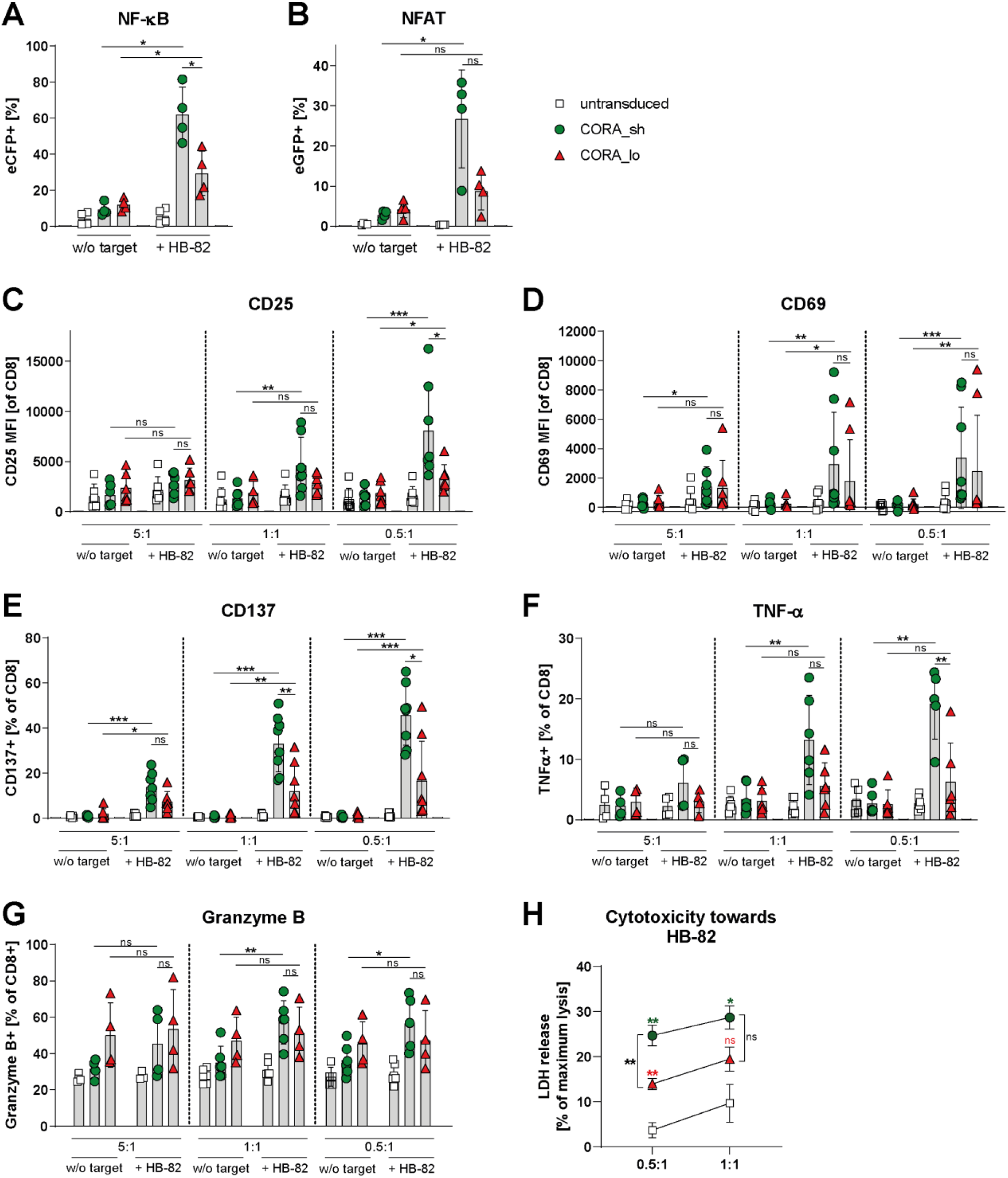
CORA_sh-Ts mediate effective and target-specific T-cell signaling, activation, cytokine expression and cytotoxicity. CORA receptors with either a short (CORA_sh) or long (CORA_lo) spacer domain were transduced into **(A, B)** Jurkat-based reporter cells or **(C-H)** primary CD8^+^ T cells isolated from healthy donors. Respective untransduced cells served as controls. **(A, B)** After cultivation of transduced reporter cells without (w/o) target cells or with HB-82 cells (anti-HLA-A*02) in an effector-to-target (E:T) ratio of 1:1 for 24 h, transcription factor activity was determined by evaluation of **(A)** NF-κB-induced enhanced cyan fluorescent protein (eCFP) or **(B)** NFAT-induced enhanced green fluorescent protein (eGFP) reporter expression by flow cytometry (n=4). **(C-H)** After co-cultivation of transduced CD8^+^ T cells with HB-82 cells in the indicated E:T ratios for 48 h, expression of **(C-E)** activation markers (n=6-8) and **(F, G)** intracellular cytokines (n=4-6) was assessed by flow cytometry. **(H)** Cytotoxicity by CORA-Ts was assessed by lactate dehydrogenase (LDH) assay. Data are shown as mean±SD (n=5). Green and red asterisks indicate comparisons of CORA_sh-and CORA_lo-Ts, respectively, with untransduced T cells. **(A-G)** Data are shown as scattered dot plot with mean±SD, whereby each symbol represents an independent donor. Statistical analysis was performed by using Mann-Whitney test. ns: not significant, *p≤0.05, **p≤0.01, ***p≤0.001.

### Modification of the CORA_sh receptor prevents activation and proliferation of CD8^+^ T cells

Application of CORA-Ts based on HLA-A*02 as recognition domain to eliminate anti-HLA-A*02 DSA-producing B cells in HLA-A*02-negative patients could potentially induce an additional sensitization of the patient’s T cells towards the HLA-A*02 within the CORA receptor. To prevent potential sensitization, we generated a CORA_sh receptor variant modified to abrogate binding of CD8^+^ T cells (CORA_sh_mod). For that, we exchanged two amino acids in the HLA-A*02 α3-domain (D227K and T228A) described to markedly reduce activation of CD8^+^ T cells by abrogation of CD8 binding to HLA-A*02 (*13*). To confirm correct assembly of the modified HLA-A*02 α-chain with cell-endogenous β2m and peptide, the CORA_sh_mod receptor was transduced into SPI-801 cells. All evaluated parameters, including detection of the correctly assembled HLA-A*02 complex on CORA_sh_mod^+^ SPI-801 cells, as well as recognition of the receptor by anti-HLA-A*02 DSAs from patient samples or by anti-HLA-A*02 antibodies from hybridoma cell supernatants (fig. S4), revealed similar results when compared to CORA_sh^+^ SPI-801 cells (fig. 1, 2, S1, S2). Thus, modification of the HLA-A*02 domain to abrogate T-cell binding did not influence expression and folding of the HLA molecule.

To prove absence of T-cell binding to CORA_sh_mod receptors, CORA_sh^+^ and CORA_sh_mod^+^ SPI-801 cells were co-cultured with CD8^+^ T cells isolated from HLA-A*02-positive healthy individuals to evaluate the T-cell response upon recognition of foreign peptides presented in the HLA-A*02 complex of CORA receptor. Following co-culture with CORA_sh^+^ SPI-801 cells for seven days, a significantly increased expression of CD25, CD69 and CD137 as markers for T-cell activation, as well as T-cell proliferation was observed (fig. 4A-E). In contrast, the same CD8^+^ T cells were not activated and did not proliferate when co-cultured with CORA_sh_mod^+^ SPI-801 cells indicating recognition by CD8^+^ T cells was prevented by the modification. Additionally, to evaluate expansion of peptide-specific T cells, CD8^+^ T cells isolated from HLA-A*02-positive healthy individuals were co-cultured with the same CORA_sh^+^ or CORA_sh_mod^+^ SPI-801 cells that were loaded with the HLA-A*02-restricted CMV pp65-derived NLV peptide before. T cells specific for the HLA-A*02/pp65_NLV_ complex specifically expanded to a frequency of 3.9% following co-culture with pp65-loaded CORA_sh SPI-801 cells compared with a frequency of 0.7% following co-culture with unloaded CORA_sh SPI-801 cells as detected by multimer staining (fig. 4F,G). In contrast, co-cultivation with NLV-loaded CORA_sh_mod^+^ SPI-801 did not increase the frequency of HLA-A*02/pp65NLV-specific T cells indicating an impaired binding to the modified CORA receptor/pp65_NLV_ complex by T cells.

Thus, application of CORA_sh_mod-Ts is expected to preclude sensitization of T cells of potential transplant and CORA-T cell therapy recipients towards foreign components of the HLA/peptide complex of the CORA receptor.

**Fig. 4:**
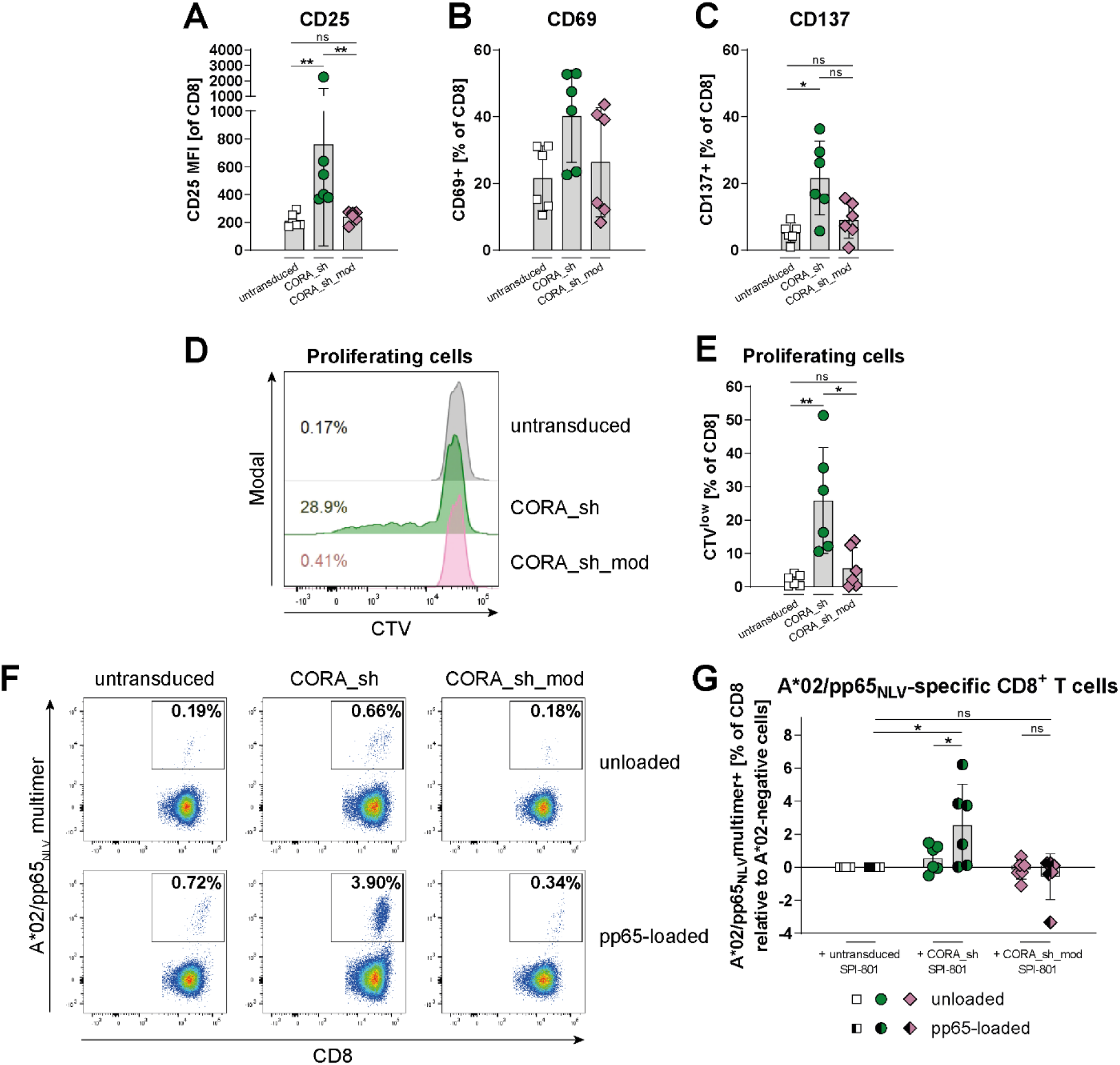
Modification of the CORA_sh receptor prevents activation and proliferation of CD8^+^ T cells. CORA_sh receptors comprising either a truncated wildtype or a modified (D227K, T228A; CORA_sh_mod) HLA-A*02 molecule were transduced into SPI-801 cells, whereby respective HLA-A*02 domains are expected to dimerize with β_2_m and to be loaded with SPI-801-derived peptides. CD8^+^ T cells were isolated from healthy, HLA-A*02^+^ donors and co-cultured with untransduced or transduced SPI-801 cells in an E:T ratio of 1:1 for 7 days. **(A-C)** Expression of activation markers and **(D, E)** proliferation of CD8+ T cells as **(D)** representative histograms or **(E)** mean±SD was assessed by flow cytometry. **(F, G)** Before co-culture with CD8^+^ T cells, transduced SPI-801 cells were exogenously loaded with the HLA-A*02-restricted and CMVpp65-derived peptide NLVPMVATV. After co-culture, the frequency of expanded HLA-A*02/pp65_NLV_-specific T cells was assessed by multimer staining using flow cytometry and is shown as **(F)** representative dot plots or **(G)** relative values, whereby respective frequencies of HLA-A*02/pp65_NLV_-specific T cells present in co-cultures with untransduced SPI-801 cells were subtracted from all values. **(A-C, E, G)** Data are shown as scattered dot plot with mean±SD, whereby each symbol represents an independent donor (n=6). Statistical analysis was performed by using **(A-C, E)** Mann-Whitney test and **(G)** Wilcoxon matched-pairs signed rank test. ns: not significant, *p≤0.05, **p≤0.01.

### CORA_sh-and CORA_sh_mod-Ts exhibit effective and target-specific T-cell signaling, activation and cytokine release

Both, the CORA_sh and the CORA_sh_mod receptor, were then evaluated for specificity and effector functionality in detail using target-positive and -negative hybridoma cell lines. In the reporter assay, significant activation of NF-κB and NFAT was induced by CORA_sh and CORA_sh_mod receptors following target recognition on anti-HLA-A*02 HB-82 and on anti-HLA-A/B/C HB-95 cells (Fig 5A,B). In that, CORA_sh_mod^+^ reporter cells increased transcription factor activation in the same extent as CORA_sh^+^ reporter cells. Importantly NF-κB and NFAT were not activated following co-culture with target-negative TÜ165 hybridoma cells proving specificity of both CORA receptors. The CORA receptors were then transduced into primary CD8^+^ T cells, which, after enrichment via EGFRt, exhibited expression of HLA-A*02 complex on 76-85% of cells assessed by either anti-HLA-A*02 antibody or HB-82 supernatant staining in flow cytometry (fig. S5A,B,D,E). However, the MFIs of both antibody stainings tended to be lower for CORA_sh_mod-Ts (2800 and 2900) when compared with CORA_sh-Ts (3400 and 4400) indicating a slightly, but not-significantly reduced receptor expression per cell (fig. S5C,F). In line with the target-specific induction of transcription factor activation (fig. 5A,B), primary CORA_sh-and CORA_sh_mod-Ts induced expression and release of effector molecules and pro-inflammatory cytokines (e.g. Granzyme B, IFN-γ, Perforin and Granulysin) and T-cell activation (indicated by upregulation of CD25, CD137) following co-culture with HB-82 and HB-95 cells, but not with TÜ165 cells (fig. 5C-E, S5G,H). Interestingly, despite similar frequencies and MFIs of BCRs on HB-82 and HB-95 cells (fig. 2B, S2A), CORA_sh and CORA_sh_mod-Ts exhibited an increased cytokine release and activation towards HB-82 when compared to HB-95 cells. To decipher reactivity of CORA-Ts towards W6/32 receptors, which is the specificity of BCRs on HB-95 cells, we utilized SPI-801 cells transduced with a membrane-bound W6/32 single chain variable fragment (scFv; W6/32 SPI-801) as additional target-positive cells and respective untransduced SPI-801 cells as negative controls to confirm anti-HLA receptor-specific T-cell functionality of CORA-Ts in detail. Upon specific recognition of expressed W6/32 scFv, transcription factor activity, release of effector molecules and T-cell activation were induced by CORA_sh and CORA_sh_mod-Ts in a significant and comparable extent, whereby both CORA-T products did not react towards SPI-801 cells (fig. 5F-J).

Taken together, CORA_sh and CORA_sh_mod-Ts exhibit distinct target specificity for anti-HLA-A*02 target cells and specifically induce multiple effector functions, whereby the amino acids substitutions for abrogation of T-cell sensitization in the CORA_sh_mod receptor did not influence target functionality.

**Fig. 5:**
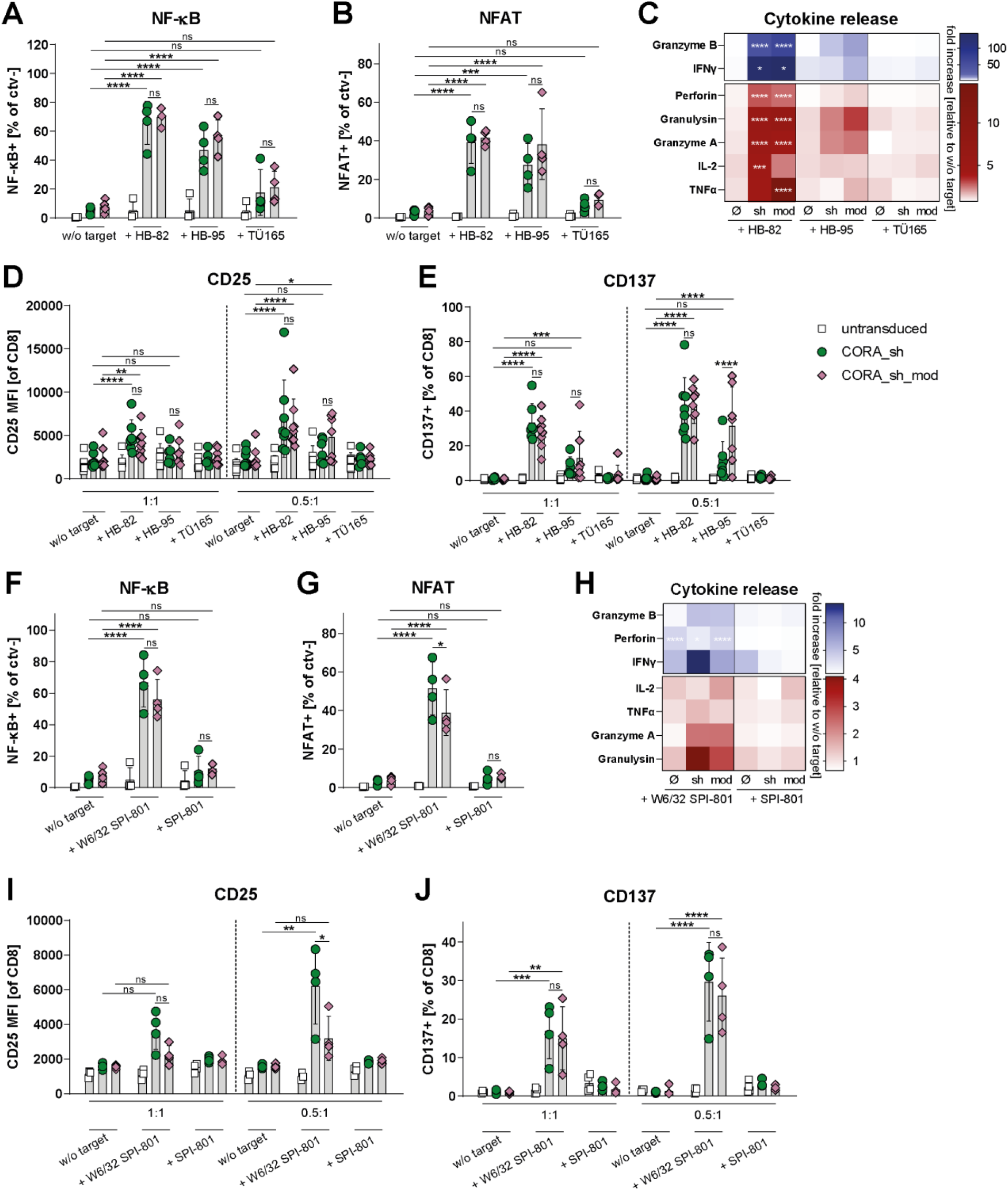
CORA_sh-and CORA_sh_mod-Ts exhibit effective and target-specific T-cell signaling, activation and cytokine release. CORA_sh receptors comprising either a truncated wildtype or a modified (CORA_sh_mod) HLA-A*02 molecule were transduced into **(A, B, F, G)** Jurkat-based reporter cells or **(C-E, H-J)** primary CD8^+^ T cells. Respective untransduced cells served as controls. **(A, B, F, G)** After cultivation of transduced reporter cells without (w/o) target cells or with the indicated target cells in an effector-to-target (E:T) ratio of 1:1 for 24 h **(A, F)** NF-κB-induced eCFP or **(B, G)** NFAT-induced eGFP reporter expression was evaluated by flow cytometry (n=4). **(C, H)** Transduced and untransduced (ø) T cells were co-cultured with the indicated target cells in an E:T ratio of 1:1 for 48 h. Release of soluble mediators into the supernatant was assessed by LEGENDplex. Fold increase to respective T cells cultures w/o target is shown as mean. (n=4-8). **(D, E, I, J)** CORA-Ts were co-cultured with target cells in the indicated E:T ratios for 48 h. Expression of activation markers was evaluated by flow cytometry. Data are shown as scattered dot plot with mean±SD, whereby each symbol represents an independent donor (n=4-10). **(A-J)** Statistical analysis was performed by using Two-Way ANOVA with Tukey’s multiple comparisons test. ns: not significant, *p≤0.05, **p≤0.01, ***p≤0.001, ****p≤0.0001.

### CORA_sh-and CORA_sh_mod-Ts mediate target-specific cytotoxicity resulting in effective reduction of anti-HLA antibody release

The cytotoxic capacity of CORA_sh- and CORA_sh_mod-Ts towards anti-HLA target cells was evaluated by co-cultivation with hybridoma cells for 48 h, followed by either analysis of living (7AAD^-^) target cells via flow cytometry or LDH assay using the co-culture supernatant. In these assays, both CORA-Ts significantly reduced viability of HB-82 and HB-95 cells, whereby CORA_sh and CORA_sh_mod-Ts exhibited an equal killing ability (fig. 6A,B,D,E). The elimination of HB-82 was higher when compared with HB-95 cells. In contrast, in line with absent induced T-cell activation and cytokine production, CORA_sh and CORA_sh_mod-Ts did not mediate cytotoxicity towards TÜ165 cells proving their specificity for anti-HLA-A*02 BCRs (fig. 6C,F). To confirm the receptor-specific functionality, W6/32 SPI-801 and corresponding untransduced SPI-801 cells were used as additional target cells (fig. 6G,H). The LDH release into the co-culture supernatant as indicator for target cell death was significantly increased in co-cultures of CORA_sh- or CORA_sh_mod-Ts with W6/32 SPI-801 cells when compared with co-cultures of untransduced T cells with W6/32 SPI-801 cells. In that, CORA_sh-Ts showed a slightly increased cytotoxic capacity when compared with CORA_sh_mod-Ts. Neither of the CORA-T products mediated elimination of untransduced SPI-801 cells confirming the W6/32 scFv as recognized target molecule.

Importantly, elimination of HB-82 cells by CORA_sh- and CORA_sh_mod-Ts lead to a significantly reduced release of anti-HLA-A*02 antibody into the co-culture supernatant as assessed by Luminex (fig. 6I). This was especially profound following an exchange of culture media after 24 h, after which anti-HLA-A*02 concentrations in supernatants of HB-82 cultured alone or together with untransduced T cells quickly increased within hours of cultivation, whereas the increase was drastically and significantly reduced in co-cultures of HB-82 cells with either CORA_sh- or CORA_sh_mod-Ts.

Thus, CORA_sh and CORA_sh_mod-Ts specifically and efficiently eliminate anti-HLA-A*02 target cells resulting in an abrogation of anti-HLA-A*02 antibody release by these cells.

**Fig. 6:**
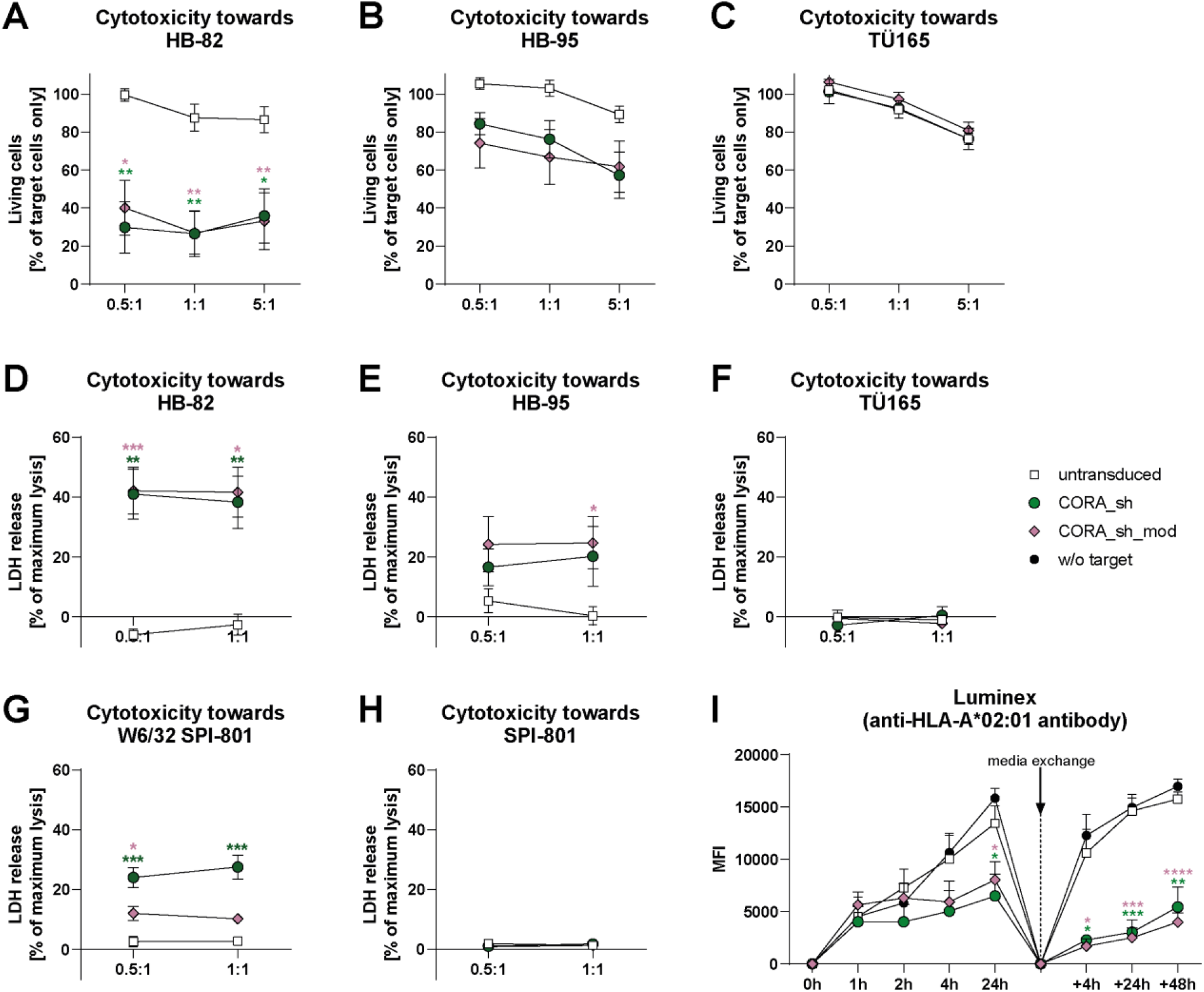
CORA_sh- and CORA_sh_mod-Ts mediate target-specific cytotoxicity resulting in effective reduction of anti-HLA antibody release. CORA_sh receptors comprising either a truncated wildtype or a modified (CORA_sh_mod) HLA-A*02 molecule were transduced into primary CD8^+^ T cells. **(A-H)** Generated CORA-Ts were co-cultured with indicated target cells in the indicated E:T ratios for 48 h. Respective co-cultures with untransduced cells served as controls. **(A-C)** Prior to the co-culture, target cells were labelled with CTV to determine the frequency of living target cells (CTV^+^7-AAD^-^) after co-culture (n=6-7). **(D-H)** Co-culture supernatants were evaluated for levels of lactate dehydrogenase (LDH) as indicator for T-cell mediated cytotoxicity (n=4-7). **(I)** Generated CORA-Ts were co-cultured with HB-82 cells in an E:T ratio of 5:1. At the indicated time points, supernatants were analyzed for released anti-HLA-A*02 antibody by Luminex. After 24 h, co-culture media were replaced by fresh medium, after which further measurements were performed at indicated time points (“+”). Data are shown as mean+SEM (n=5). **(A-H)** Data are shown as mean±SEM. Statistical analysis was performed by using Two-Way ANOVA with Dunnett’s multiple comparisons test. **(A-H)** Significances are shown in comparison with respective untransduced T cells cultured in the same E:T ratio. **(I)** Significances are shown in comparison with HB-82 cultured alone and evaluated at the respective same time points. *p≤0.05, **p≤0.01, ***p≤0.001, ****p≤0.0001.

### Tacrolimus-resistant CORA-Ts sustain target-specific effector functions in presence of immunosuppression

As therapy to prevent organ rejection, most SOT patients are continuously administered immunosuppressive medication, e.g. the calcineurin inhibitor tacrolimus. To overcome a reduction of CORA-T functionality under immunosuppressive conditions, they were endowed with resistance towards tacrolimus treatment as previously demonstrated for CMV-specific T cells (*14*). For that, besides transduction with the CORA_sh receptor, a CRISPR/Cas9-mediated knockout of the tacrolimus-binding protein gene *FKBP12* achieving CRISPR efficiencies of 80% was implemented in the CORA-T manufacturing process (fig. 7A). Functionality of generated tacrolimus-resistant FKBP^ko^-CORA_sh-Ts in comparison with unmodified CORA_sh-Ts was tested by co-cultivation with HB-82 cells with or without addition of tacrolimus in a clinically relevant dose (*15*). Upregulation of CD25 and CD137 on CORA_sh-Ts following co-culture with HB-82 cells as markers for T-cell activation was significantly reduced in presence of tacrolimus when compared with respective co-cultures without tacrolimus addition (fig. 7B,C). In contrast, activation of FKBP^ko^-CORA_sh-Ts following recognition of HB-82 cells was not altered by the presence of tacrolimus. Cytotoxicity of CORA_sh-Ts towards HB-82 cells was not reduced after co-cultivation for 48 h in presence of tacrolimus, which is why they were re-stimulated with new HB-82 cells (fig. 7D). Towards newly-added HB-82 cells, CORA_sh-Ts tended to exhibit a reduced cytotoxicity in presence of tacrolimus, whereas FKBP^ko^-CORA_sh-Ts mediated equally high cytotoxicity after both stimulations irrespective of immunosuppressive treatment. In the same re-stimulation co-culture with HB-82 cells, T-cell proliferation measurements by CTV dilution assay revealed a significantly diminished proliferation of CORA_sh-but not of FKBP^ko^-CORA_sh-Ts in presence of tacrolimus (fig. 7E,F). In line, the release of several pro-inflammatory cytokines and effector molecules (e.g. TNF-α, IFN-γ and granulysin) in response to HB-82 recognition was significantly reduced by tacrolimus addition to CORA_sh-T, but not to FKBP^ko^-CORA_sh-T co-cultures [Fig 7G].

Thus, CRISPR/Cas9-mediated knockout of the tacrolimus-binding protein FKBP12 endowed CORA-Ts with the ability to mediate potent effector functions irrespective of the presence of tacrolimus.

Our results demonstrate that by engineering of T cells with an innovative CORA receptor, they are able to specifically recognize and eliminate distinct anti-HLA B cells, having the potential to selectively prevent the formation of anti-HLA antibodies even under immunosuppressive conditions. This suggests CORA-Ts as a potent novel approach to specifically eliminate alloreactive anti-donor-HLA B cells to combat AMR and to improve long-term graft survival in SOT patients while preserving their overall B-cell immunity.

**Fig. 7:**
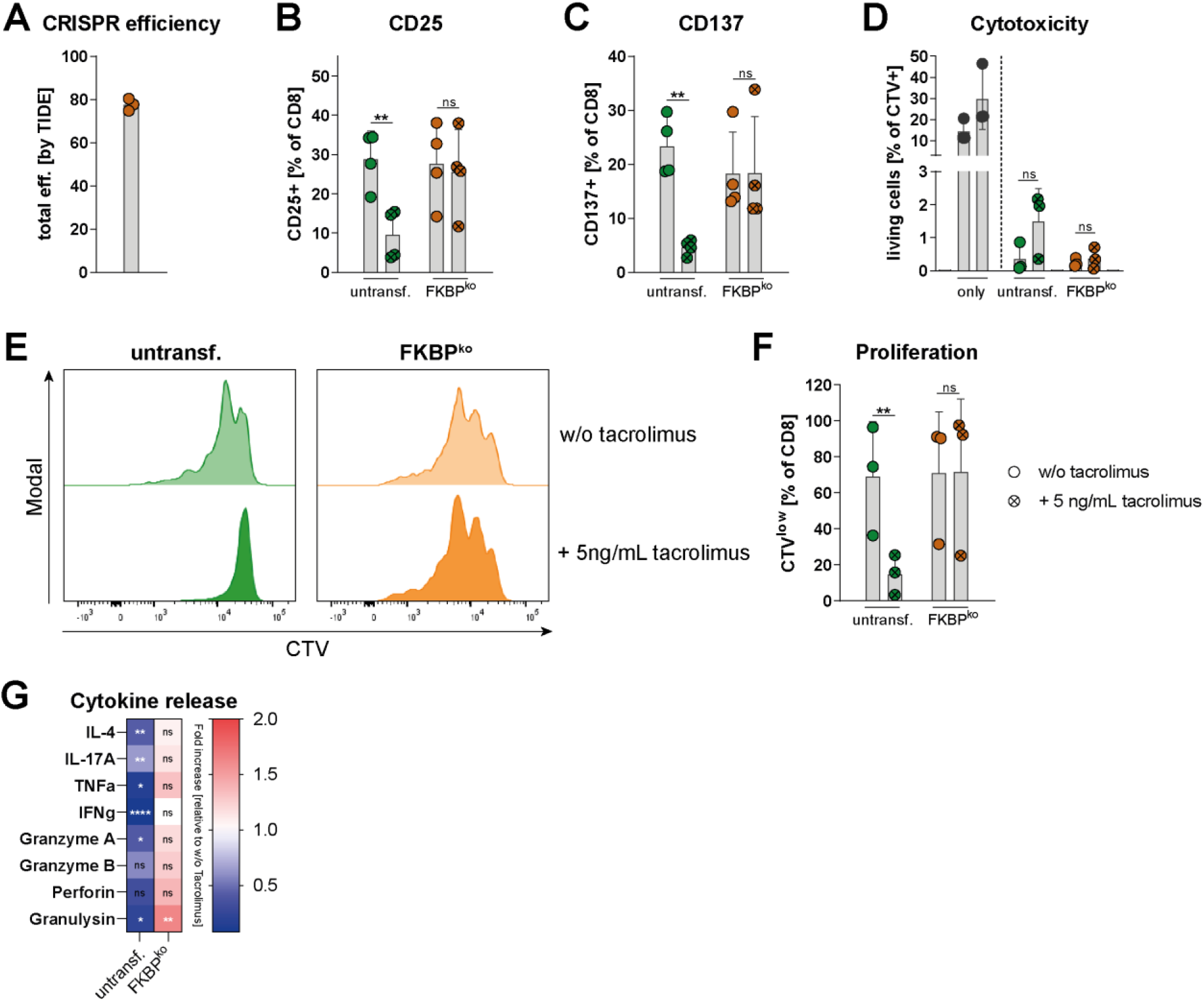
Tacrolimus-resistant CORA-Ts sustain target-specific effector functions in presence of immunosuppression. CORA_sh receptors were transduced into CD8^+^ T cells, followed by transfection with FKBP12-targeting RNP complex (FKBP^ko^). Respective untransfected CORA-Ts served as controls (untransf.). **(A)** CRISPR efficiency was assessed by analysis of sequencing results by TIDE (*16*). **(B-C)** CORA-Ts were co-cultured with HB-82 cells for 2 days in an E:T ratio of 1:1 in presence or absence (w/o) of 5 ng/mL tacrolimus, after which expression of activation markers was evaluated by flow cytometry (n=4). **(D-G)** Prior to the co-culture, CORA-Ts were labelled with CTV. After 2 days of co-culture, CORA-Ts were re-stimulated with the same number of HB-82 cells (n=3). After 5-7 days of co-culture, **(D)** the frequency of living target cells (CTV^-^7-AAD^-^) was assessed by flow cytometry. HB-82 cells cultured alone (only) for 5-7 or 3-5 days, respectively, are shown as first and second bar. **(E, F)** Proliferation of CORA-Ts was assessed via CTV dilution assay by flow cytometry and is shown as **(E)** representative histograms or **(F)** mean. **(G)** Release of soluble mediators into the supernatant was assessed by LEGENDplex. Fold increase to respective co-cultures in absence of tacrolimus is shown as mean. Statistical analysis was performed by using Two-Way ANOVA with Šídák’s multiple comparisons test. **(B-D, F)** Data are shown as scattered dot plot with mean+SD, whereby each symbol represents an independent donor. Statistical analysis was performed by using Two-Way ANOVA with Tukey’s multiple comparisons test. ns: not significant, *p≤0.05, **p≤0.01, ****p≤0.0001.

## Discussion

In the present study, we developed a cell therapeutic approach to specifically eliminate anti-donor-HLA B cells and thereby abrogate release of anti-HLA DSAs responsible for AMR, while preserving B cells with different specificities. To redirect autologous T cells against distinct anti-HLA B cells, we designed a CORA receptor that is composed of an HLA molecule of interest fused to 4-1BB and CD3ζ T-cell signaling domains commonly used for CARs (*17*). As proof-of-concept, a truncated HLA-A*02 α-chain was encoded in the CORA receptor and shown to be assembled with β_2_m and peptides by the cell-endogenous machinery. A short spacer domain between HLA α-chain and transmembrane domain in the CORA_sh receptor enabled superior HLA-complex expression on all transduced cells when compared with a long spacer domain variant. In line, recognition of anti-HLA-A*02 BCRs on the surface of HB-82 cells (anti-HLA-A*02), and effector functionality including cytotoxicity towards the B-cell line was improved for CORA_sh-Ts when compared with CORA_lo-Ts, matching to studies in CAR-T cells that described a correlation between effector functionality and density of CAR expression (*18*).

CORA_sh-Ts were moreover proven to be highly specific by utilizing additional hybridoma cells and artificial target cells with W6/32 specificity, revealing potent and anti-HLA-A*02-specific induction of T-cell signaling and activation, release of cytokines and effector molecules, as well as cytotoxicity. As main aim in the approach to combat AMR, they moreover efficiently reduced the release of anti-HLA-A*02 antibodies. In contrast, CORA_sh-Ts did not react towards target-negative cells including the anti-HLA-B*35 B-cell line TÜ165, proving their selectiveness. Thus, as opposed to currently available B-cell depletion or immunosuppression protocols (*3*), CORA-T-cell therapy would potentially preserve the general humoral immunity for defense against pathogens and maintain supportive B_reg_ subpopulations beneficial for allograft tolerance.

CORA_sh-Ts reacted towards all anti-HLA-A*02 target cells utilized in the present study, yet to a different extent. Although transcription factor activity and T-cell activation were induced in CORA_sh-Ts in a similar extent following co-culture with HB-82, HB-95 and W6/32 SPI-801 cells, markedly less cytokines and effector molecules were released by CORA_sh-Ts in co-cultures with HB-95 and W6/32 SPI-801 compared with HB-82 cells.

Moreover, CORA_sh-Ts exhibited reduced cytotoxicity towards HB-95 and W6/32 SPI-801 cells, both of which share W6/32 specificity, compared with HB-82 based on BB7.2. This is in line with the lower frequency of HB-95 cells that could be stained with eukaryotic HLA-A*02 multimer when compared with HB-82 cells, although both, frequency and MFI, of anti-mIgG staining of BCRs in flow cytometry were comparable between HB-82 and HB-95 cells. Nevertheless, CORA_sh-Ts mediated effective cytotoxicity towards all anti-HLA-A*02 target cells within 48 h. Hence, they are expected to achieve a complete clearance over time due to the usage of a 4-1BB-based receptor design described to mediate long-term persistence of CAR-T cells (*19*). In the future, *in vivo* models will be required to evaluate long-term target elimination and persistence of CORA-T cells.

As indicator that CORA-Ts should be capable to target also alloreactive naïve and memory anti-HLA-A*02 B cells targeting various epitopes in transplant patients, anti-HLA-A*02 DSAs, i.e. soluble BCRs, present in the serum of HLA-A*02-negative transplant patients were shown to be recognized by the CORA_sh receptor.

As potential drawback of a CORA-Ts as new therapy approach, whereas alloreactive naïve and memory B cells are characterized by anti-HLA BCRs, plasma cells express little or no surface BCR. Thus, equally to current B-cell depletion protocols targeting CD20, plasma cells are not expected to be efficiently eliminated. A promising approach could be the combination with the proteasome inhibitor bortezomib to target plasma cells and prevent reconstitution of anti-donor-HLA plasma cells from memory B cells by application of CORA-Ts. However, all plasma cells including protective virus-specific cells would be depleted by bortezomib treatment. Otherwise, pre-treating nonsensitized patients on the waiting list with CORA-Ts to prevent development of AMR by depletion of naïve HLA-specific B cells would circumvent formation of corresponding plasma cells but retain the general B-cell immunity.

As additional combination, plasmapheresis could prevent potential masking and reduction in functionality of CORA-Ts by DSAs present in the blood of transplant patients. This is contradicted by data from a cell therapy approach targeting autoreactive B cells via their BCRs in the context of pemphigus vulgaris (PV) (*20*). Here, soluble autoantibodies interacting with the engineered T-cell receptor were shown to induce low-level proliferation of the engineered T-cells expected to increase their efficacy and persistence (*21*). Functionality of these chimeric autoantigen receptor (CAAR) T cells could impressively be demonstrated using different *in vitro* and *in vivo* models, leading to their evaluation in an ongoing, currently recruiting clinical study (NCT04422912). One similar approach was developed to target B cells releasing neutralizing antibodies against the replacement factor VIII in hemophilia A patients and such B-cell antibody receptor (BAR)-engineered T cells significantly reduced the anti-factor VIII antibody formation *in vivo* (*22*). Thus, these studies proof general suitability of a cell therapy approach targeting B cell populations via their BCRs.

The CORA_sh receptor was further modified by the exchange of two amino acids (D227K and T228A) in the HLA-A*02 α_3_ domain, which did not influence overall functionality of CORA-Ts. These amino acids are known to be part of a negatively-charged α_3_-chain loop important for binding of CD8 (*23*). Purbhoo *et al*. could show that the utilized substitutions in HLA-A*02 multimers were sufficient to efficiently reduce CD8^+^ T-cell activation following multimer staining (*13*). In line, in the present study, the CORA_sh_mod receptor, in contrast to the unmodified CORA_sh receptor, did not induce activation or proliferation of CD8^+^ T cells specific for HLA-A*02 in complex with foreign, SPI-801-derived peptides or loaded with a distinct CMV pp65-derived peptide, most likely due to the described abrogation of CD8 binding. Thus, CORA_sh_mod-Ts are not expected to interact with alloreactive anti-donor-HLA T cells in the patient, which increases safety of the approach. Application of CORA_sh-Ts, due to the foreign HLA expressed within the CORA receptor, could potentially induce an additional sensitization towards donor-HLA molecules and exacerbation of TMR towards the allograft. On the other side, CORA_sh-Ts might have the potential to eliminate alloreactive anti-donor-HLA T cells upon binding of HLA-A*02-specific T cells to the HLA-A*02 domain of the CORA receptor. Thus, it will be interesting to study the potential of CORA_sh-Ts as tool to not only combat AMR but also TMR in a future study. As approach to eliminate T cells with distinct peptide specificity in context of autoimmune diseases, Yi *et al*. reported generation of T cells engineered with a peptide-HLA class II CARs and showed their ability to specifically deplete the respective peptide-reactive CD4^+^ T cells in a mouse model (*24*). Moreover, Quach *et al*. used a β_2_m CAR that was able to complex with cell-endogenous HLA α-chains to protect virus-specific T cells from elimination by alloreactive PBMCs *in vitro* as approach for an off-the-shelf cell therapy (*25*).

As further modification, CRISPR/Cas9-mediated k.o. of the corresponding binding protein gene *FKBP12* endowed CORA-Ts with potent protection from tacrolimus treatment. Calcineurin inhibitors, such as tacrolimus, are currently the most effective and therefore a widely-used class of maintenance immunosuppression following SOT (*14*), since they potently reduce the activation, maturation and cytokine secretion by T cells (*26*). Thus, tacrolimus-resistant CORA-Ts are a promising approach with expected high functionality even under immunosuppressive conditions. Especially CORA_sh-Ts to potentially target TMR would benefit from combination with tacrolimus treatment to receive an advantage over the suppressed alloreactive T cells.

We here present a promising new tool to selectively eliminate alloreactive anti-HLA B cells responsible for AMR even under immunosuppressive conditions while preserving the protective B-cell immunity. By using CORA-Ts targeting anti-HLA-A*02 B cells as a proof-of-concept, a smooth development of CORA-Ts based on further HLA class I and class II alleles broadens the applicability to a larger cohort of transplant patients in an individual and patient-tailored manner.

## Material and Methods

### Human sample materials

All experiments with primary cells were performed using residual blood samples from routine platelet collection at the Institute of Transfusion Medicine and Transplant Engineering, Hannover Medical School. According to standard donation requirements, the respective donors had no signs of acute infection and no previous history of blood transfusion. Written informed consent was obtained from all donors as approved by the Ethics Committee of Hannover Medical School (2519–2014, 3639–2017). Residual frozen sera samples from seven kidney transplant recipients were evaluated, which was approved by the Ethics Committee of Hannover Medical School (8969_BO_K_2020).

### Construction of CORA vectors

To design the CORA receptors, sequences of HLA-A*02:01:01:01 exons 1-4 (GenBank no. HG794376.1) were synthesized (Thermo Fisher Scientific) and cloned into two previously described TÜ165-CAR-epHIV7 vectors by using NheI and RsrII restriction sites to, respectively, replace the signaling peptide and TÜ165 scFv (*27*). Briefly, the resulting receptors comprised the extracellular chains α_1_-α_3_ of HLA-A*02:01 fused to either a short (CORA_sh) “hinge-only” (12 aa) or a long (CORA_lo) “hinge-CH2-CH3” (229 aa) spacer domain region derived from IgG4-Fc followed by a transmembrane domain (TMD) of CD28 and the intracellular signaling domains of 4-1BB and CD3ζ (Figure 1A). EGFRt (*28*) was encoded in the same vector by using a self-cleaving T2A element and served as marker for detection and enrichment of transduced cells.

To generate CORA receptors modified to abrogate T-cell binding to the HLA-A*02 domain (CORA_sh_mod and CORA_lo_mod), two residues (D227K and T228A) in the α_3_ domain of HLA-A*02:01 described to abrogate binding of CD8 (*13*) were changed by site-directed mutagenesis (Agilent).

### Generation of lentivirus

CORA_sh, CORA_lo, CORA_sh_mod and CORA_lo_mod lentiviral particles were produced similar as described before (*27*). Briefly, 293T cells (ACC 635; DSMZ) were transfected with respective CORA-epHIV7 vectors and third-generation packaging vectors in presence of 25 µM chloroquine. Supernatants containing lentivirus were harvested after 32h and 48h and concentrated via ultracentrifugation. Titers were determined by transduction of Jurkat cells (ACC 282; DSMZ) in presence of 5 µg/ml Polybrene Infection/Transfection Reagent (Merck). After 48 h, transduction efficiencies assessed by staining of co-expressed EGFRt in flow cytometry were used to calculate virus titers.

### Cell lines

SPI-801 cells (ACC 86; DSMZ), which are HLA-negative K562 cells, were transduced with respective CORA receptors in a multiplicities of infections (MOI) of 1 and 5 μg/mL Polybrene (Merck). Cells harboring the receptor were enriched by using biotinylated anti-EGFR antibody and anti-biotin microbeads (Miltenyi Biotec).

The hybridoma cell lines HB-82 (anti-HLA-A*02; clone BB7.2; ATCC), HB-95 (anti-HLA class I; clone W6/32; ATCC) and TÜ165 (against HLA-B*35:01 loaded with LPPHDITPY; kindly provided by Barbara Uchanska-Ziegler (Ziegler Biosolutions, Waldshut-Tiengen, Germany (*29*))) were used as surrogates for B cells expressing anti-HLA BCRs and releasing the respective antibody. Cell culture supernatants containing respective anti-HLA antibodies were harvested from all hybridoma cells. W6/32 SPI-801 cells were generated by transduction of a CAR construct based on the scFv of the anti-HLA class I antibody W6/32 into SPI-801 cells by lentiviral transduction.

### Detection of anti-HLA-A*02 antibodies

Serum of kidney transplant recipients were evaluated for presence of anti-HLA-A*02:01 antibodies using mixed HLA antigen-charged polysterene beads (LIFECODES LifeScreen LSA test, Gen-probe-Immucor) and a multi-analyte flow array (Luminex® 200™ System) according to the manufacturer’s instructions. Murine anti-HLA-A*02:01 antibodies released into cell culture supernatants by HB-82 cells were analogously evaluated, except for the usage of an anti-mouse IgG (H+L) polyclonal F(ab’)_2_ fragment (Jackson Immunoresearch) for detection.

### Crossmatch

PBMCs or CORA-receptor transduced SPI-801 cells were incubated with serum of kidney transplant recipients with detectable anti-HLA-A*02:01 concentrations or cell culture supernatants of hybridoma cells for 30 min. Rabbit complement (BioRad) was added and incubated for 60 min. FluoroQuench Stain/Quench reagent (Thermo Fisher Scientific) was used to distinguish live cells (green) from dead cells (red) by using a Keyence BZ-8100E microscope. Crossmatch scores were determined by using values 1 (≤ 10%), 2 (10-20%), 3 (20-40%), 6 (40-80%) and 8 (80-100%) for increasing frequencies of dead cells according to standard protocols of the National Institute of Health (USA) (*11*).

### Reporter assay to determine CORA-receptor signaling

CORA receptors were transduced into a previously described Jurkat-based reporter cell line (*12*) by addition of lentiviral particles in a MOI of 1 and 5 μg/mL Polybrene (Merck) using spinoculation. After 48 h, transduced reporter cells were co-cultured with the indicated target cells previously labelled with CellTrace^TM^ violet proliferation dye (CTV; Thermo Fisher Scientific) for 24 h in an effector to target (E:T) ratio of 1:1. CORA receptor-induced signaling was assessed by detection of enhanced cyan fluorescent protein (eCFP) reporting for NF-κB and enhanced green fluorescent protein (eGFP) reporting for NFAT activation in CTV^-^ reporter cells by flow cytometry. Untransduced reporter cells were treated analogously and used as control.

### Generation of primary (tacrolimus-resistant) CORA-Ts

Primary CORA-Ts were generated as previously described (*27*). Briefly, Peripheral blood mononuclear cells (PBMCs) were isolated from residual blood samples from routine platelet collection of healthy individuals by density gradient centrifugation using Lymphosep (c.c.pro). CD8^+^ T cells were isolated (Miltenyi Biotec), activated with anti-CD3/CD28 beads (Thermo Fisher Scientific) in a ratio of 1:1, and cultured in TexMACS (Miltenyi Biotec) with 3% human serum (c.c.pro; CTL medium) supplemented with 12.5 ng/ml IL-7 and IL-15 (PeproTech). On day 1, T cells were transduced with CORA receptors by addition of lentivirus in an MOI of 3, 5 µg/mL Polybrene and spinoculation. On day 2, the anti-CD3/CD28 beads were removed. For the generation of tacrolimus-resistant CORA-Ts, beads were removed on day 3 followed by electroporation using the Human T Cell Nucleofector™ Kit (Lonza) and the Amaxa Nucleofactor^®^ 2b (Lonza, program T-023) to transfer the RNP complex of 62 pmol of Alt-R *Streptococcus pyogenes* Cas9 protein V3 precomplexed with 72 pmol duplex of FKBP12-targeting crRNA (5’-GGGCGCACCTTCCCCAAGCG-3’; sequence previously described (*14*)) and tracrRNA (all from Integrated DNA Technologies). After that, the cells were split depending on their growth. On day 8-9, transduced T cells were enriched via co-expressed EGFRt. EGFRt^+^ cells were then further expanded until day 12-16. Untransduced T cells were expanded analogously, except for the addition of lentivirus and enrichment of EGFRt^+^ cells, and served as control.

For evaluation of on-target editing efficiency of tacrolimus-resistant CORA-Ts, DNA was isolated on day 12-16 by using the DNeasy Blood and Tissue Kit (Qiagen). The FKBP12 site was amplified with the Platinum^TM^ SuperFi DNA Polymerase (Thermo Fisher Scientific) according to the manufacturer’s instructions and using the following primers: 5’-TCTGACGGGTCAGATAACACCTAG-3’ and 5’-TCTTCCGGAGGCCTGGGTTT-3’ (as previously described (*14*)). PCR products were purified (Qiagen), Sanger sequenced (Microsynth Seqlab), and CRISPR efficiency was determined by TIDE software version 3.3.0 (*16*).

### Evaluation of CORA-T functionality

CORA-T functionality was evaluated following co-culture with indicated target cells in CTL medium. For detection of anti-HLA-A*02 antibodies, samples of co-culture supernatants were taken at the indicated time points, whereby media was replaced with new CTL medium after 24 h. For evaluation of tacrolimus-resistant CORA-Ts, they were co-cultured with HB-82 cells in absence or presence of 5 ng/mL tacrolimus (Merck) in an E:T ratio of 1:1 for 48 h. Cytotoxicity, proliferation and cytokine release were assessed after 5-7 days, whereby T cells were re-stimulated with the same number of HB-82 cells after 2 days.

Abrogation of CD8 binding to the CORA_sh_mod receptor was evaluated by co-cultivation of CORA-receptor-transduced SPI-801 cells with CD8^+^ cells isolated from PBMCs of healthy HLA-A*02-positive individuals in an E:T ratio of 1:1 for 7 days.

### Flow cytometry

Antibodies used for flow cytometry are listed in table S1. To stain HLA-A*02:01, cell culture supernatant of HB-82 cells containing anti-HLA-A*02:01 antibody (mIgG2b) was utilized and detected with anti-mouse IgG secondary antibody (Jackson Immunoresearch). T-cell proliferation was assessed by labelling CORA-Ts with CTV before co-culture with target cells and gating on CTV^low^ cells. Samples were analyzed on a BD FACSCanto Flow Cytometer (Becton Dickinson). Data were analyzed using FlowJo v10.

### Multiplex cytokine analysis

To determine effector molecule concentrations in co-culture supernatants, the LEGENDplex human CD8/NK Panel (BioLegend) was performed according to the manufacturer’s instructions. Data were analyzed with LEGENDplex v8.0 software (BioLegend). Fold-increase was calculated by division by concentrations of the respective CORA-Ts cultured alone.

### Cytotoxicity

Killing capacity of CORA-Ts was assessed by co-cultivation with CTV-labelled target cells for 48 h. Afterwards, 7-AAD (BioLegend or BD) was used to discriminate dead CTV^+^ cells via flow cytometry. As alternative, cytotoxicity was determined based on the release of lactate dehydrogenase (LDH) into the co-culture supernatant measured by Cytotoxicity Detection Kit (Roche) and using a Synergy 2 Multi-Mode Microplate Reader (Biotek) to determine absorbance at a wavelength of 490 nm with a reference wavelength of 690 nm. Maximum lysis was measured by addition of 1% Triton X-100 (Merck) to control wells. LDH release by eliminated target cells in co-cultures with CORA-Ts was calculated by subtracting absorbance values measured in wells with respective target and CORA-Ts cultured alone and divided by the respective maximum lysis control of the co-culture.

### Statistical analysis

Statistical analysis was performed by Graph Pad Prism 9.5.1 using Mann-Whitney T tests or two-way ANOVA with Dunnet’s or Tukey’s multiple comparisons tests as indicated. ns: not significant, *p≤0.05; **p≤0.01; ***p≤0.001; ****p≤0.0001.

## Acknowledgments

We thank Elvira Schulde, Dörthe Rokitta and Sarina Lukis (all Institute of Transfusion Medicine and Transplant Engineering, MHH, Hannover, Germany) for their technical support. Moreover, we would like to thank Dr. Anja Battermann and Philipp Schleumann (both imusyn GmbH & Co. KG, Hannover, Germany) for advice for cloning and staining, as well as providing the eukaryotic HLA-A*02 multimer.

## Author contributions

Conceptualization: ACD, CFF, RB, BEV

Methodology: ACD, AB, MV

Investigation: ACD, AB, MV, BEV

Visualization: ACD

Funding acquisition: ACD, RB, BEV

Project administration: ACD, RB, BEV

Supervision: MH, CFF, RB, BEV

Writing – original draft: ACD, AB, BEV

Writing – review & editing: ACD, AB, MV, MH, CFF, RB, BEV

## Data and materials availability

Data are available upon reasonable request. The data sets and protocols used and/or analyzed during the current study are available from the corresponding author on reasonable request.

