## Supplementary Materials for "Depletion of alloreactive B cells by chimeric alloantigen receptor T cells with drug resistance to prevent antibody-mediated rejection in solid organ transplantation"

### Supplementary Figures

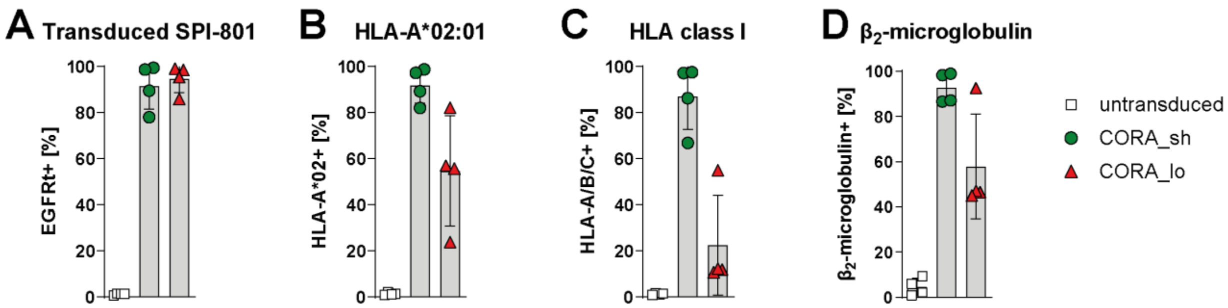


**Fig. S1: CORA receptors harbor a truncated HLA-A*02 as recognition domain.** CORA_sh and CORA_lo receptors were generated and expressed in SPI-801 cells by lentiviral transduction. Expression of **(A)** EGFRt, **(B)** HLA-A*02, **(C)** HLA class I and **(D)** $\beta_2$-microglobulin was assessed by flow cytometry. Untransduced SPI-801 cells served as control. Data are shown as scattered dot plot with mean±SD, whereby each symbol represents an independent experiment (n=4).

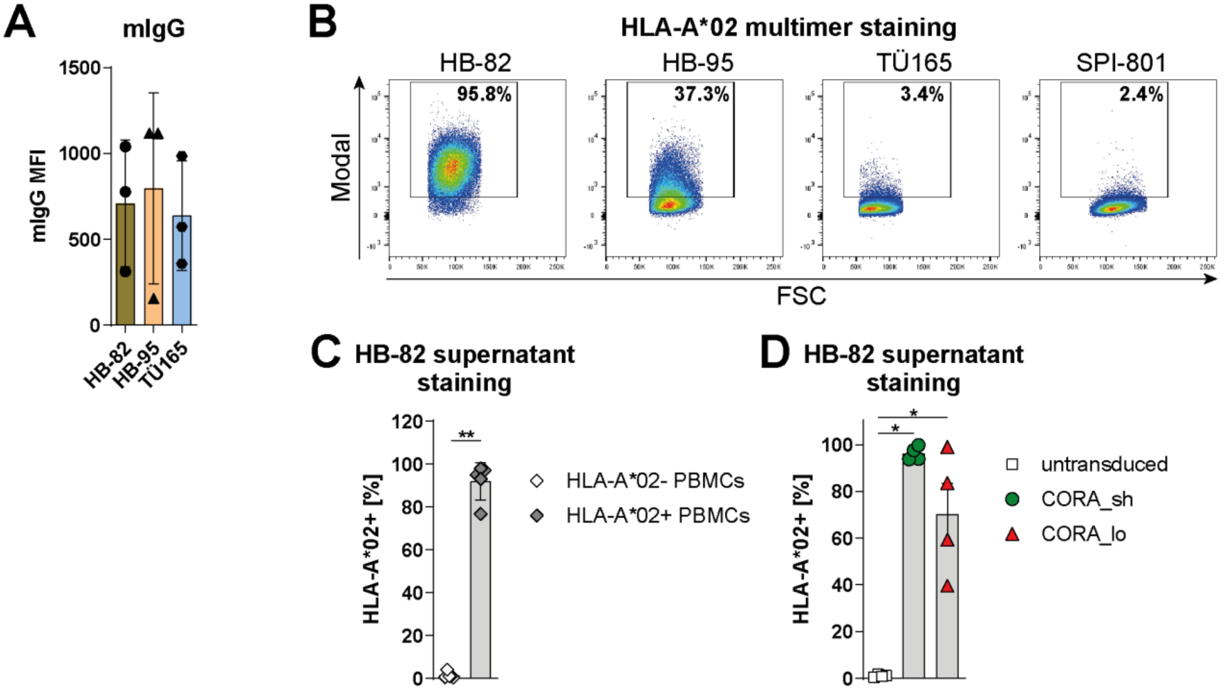


**Fig. S2: Hybridoma cells serve as model for anti-HLA-A*02 B cells releasing anti-HLA*A02 antibodies.** HB-82 (anti-HLA-A*02), HB-92 (anti-HLA-A/B/C) and TÜ165 (anti-HLA-B*35 loaded with LPPHDITPY) cells were used as model for anti-HLA-antibody-releasing B cells. **(A)** Surface expression of murine BCRs was determined by flow cytometry and usage of anti-mouse immunoglobulin G (mIgG) antibody. **(B)** Recognition of HLA-A*02 by hybridoma cells was assessed by staining with eukaryotic HLA-A*02/NLV multimer. **(C, D)** Cell culture supernatant of HB-82 cells containing anti-HLA-A*02 antibody was used to stain **(C)** HLA-A*02-negative or -positive PBMCs from healthy donors, as well as

**(D)** SPI-801 cells transduced with CORA_sh or CORA_lo receptors. **(A, C-D)** Data are shown as scattered dot plot with mean±SD, whereby each symbol represents an independent experiment (n=3-5). Statistical analysis was performed by using Mann-Whitney test. *p≤0.05, **p≤0.01.

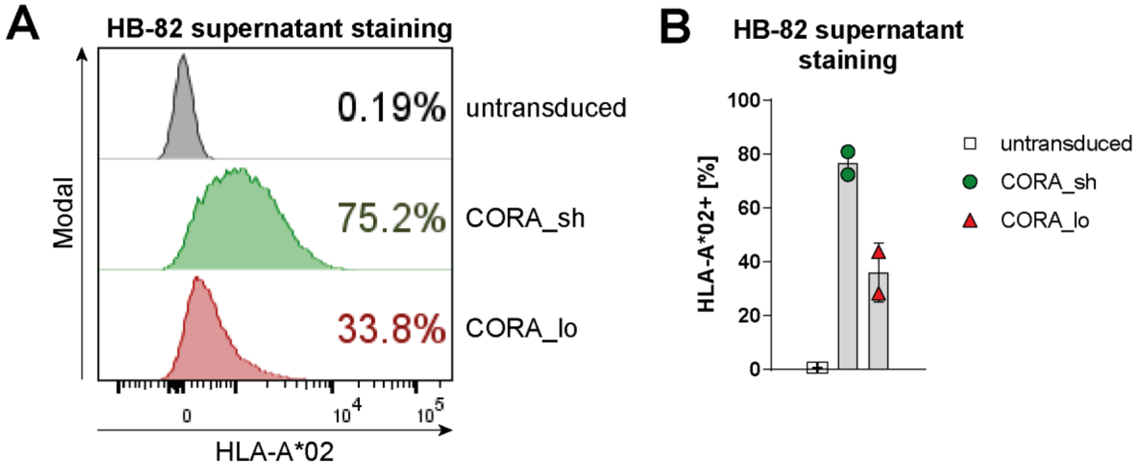


**Fig. S3: CORA receptors with a truncated HLA-A*02 molecule as recognition domain are detectable on the cell surface after transduction into primary CD8⁺ T cells.** CORA receptors with either a short (CORA_sh) or long (CORA_lo) spacer domain were transduced into primary CD8⁺ T cells isolated from healthy donors. CORA receptor⁺ cells were enriched using EGFRt as selection marker. After manufacturing, HLA-A*02 expression on the cell surface was assessed by flow cytometry and staining with cell culture supernatant of HB-82 cells containing anti-HLA-A*02 antibody. Data are shown as **(A)** representative histograms or **(B)** scattered dot plot with mean±SD, whereby each symbol represents an independent donor (n=2).

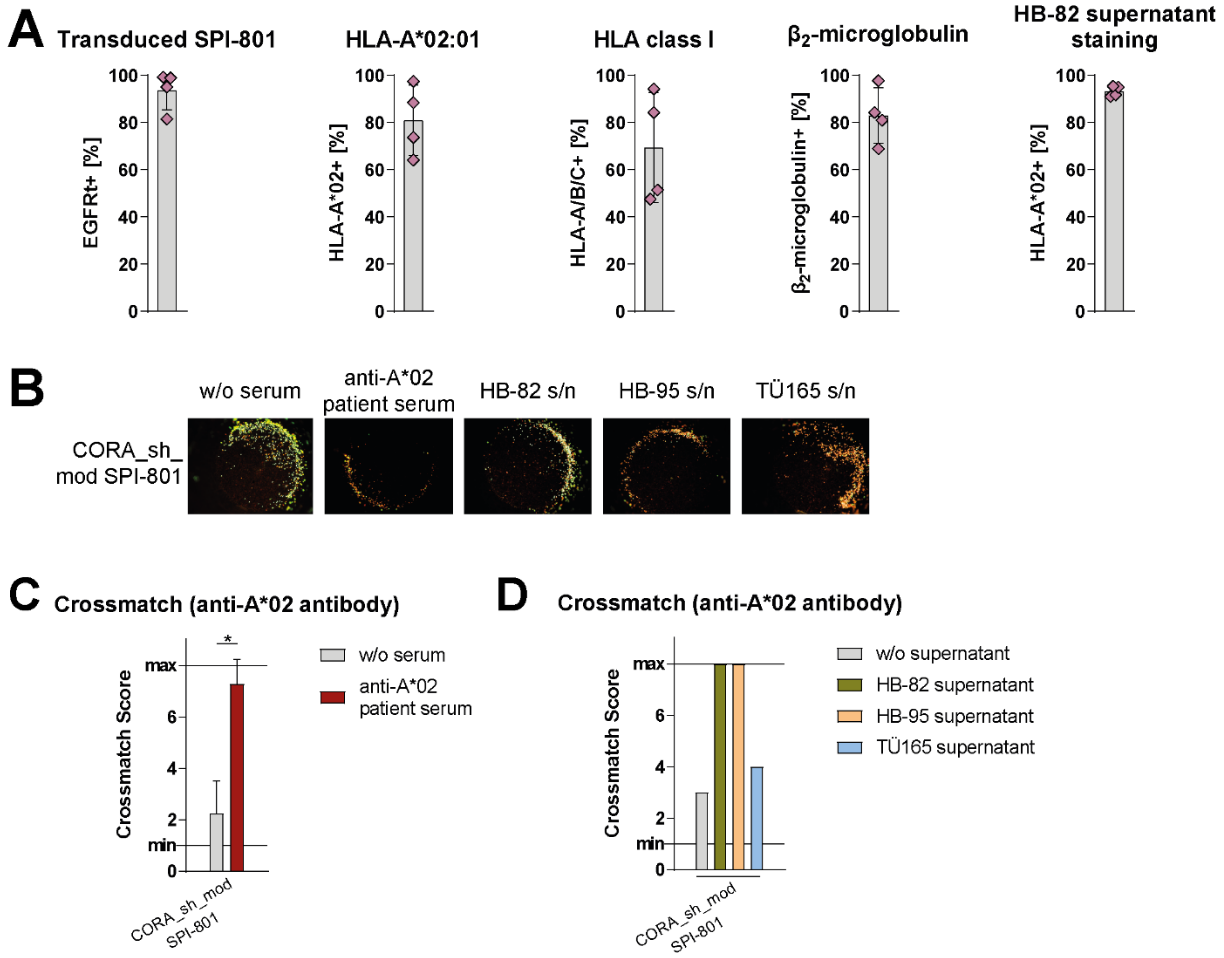


**Fig. S4: Modification of the HLA-A*02 component of CORA receptors to abrogate CD8 binding does not interfere with HLA complex association nor recognition by antibodies.** CORA receptors comprising a modified truncated HLA-A*02 molecule (D227K, T228A) and a short spacer domain (CORA_sh_mod) were transduced into SPI-801 cells. **(A)** Expression of EGFRt, HLA-A*02, HLA class I and $\beta_2$-microglobulin was assessed by flow cytometry. Moreover, cell culture supernatant of HB-82 cells containing anti-HLA-A*02 antibody was used to stain CORA_sh_mod$^+$ SPI-801 cells (right graph). Data are shown as scattered dot plot with mean±SD, whereby each symbol represents an independent experiment (n=4). **(B-D)** Binding of anti-HLA antibodies present in **(B, C)** the serum of kidney transplant recipients or **(B, D)** the supernatant (s/n) of hybridoma cells to CORA_sh_mod$^+$ SPI-801 cells was assessed by their ability to mediate complement-dependent cytotoxicity (CDC) in crossmatch assays. **(B)** Representative pictures and **(C, D)** crossmatch scores indicate CDC based on evaluation of viable cells (green) versus dead cells (red) after complement addition. Respective cells incubated without (w/o) supernatant served as viable controls. **(C)** Data are shown as mean+SD (n=4-10). Statistical analysis was performed by using Mann-Whitney test. *$p \leq 0.05$.

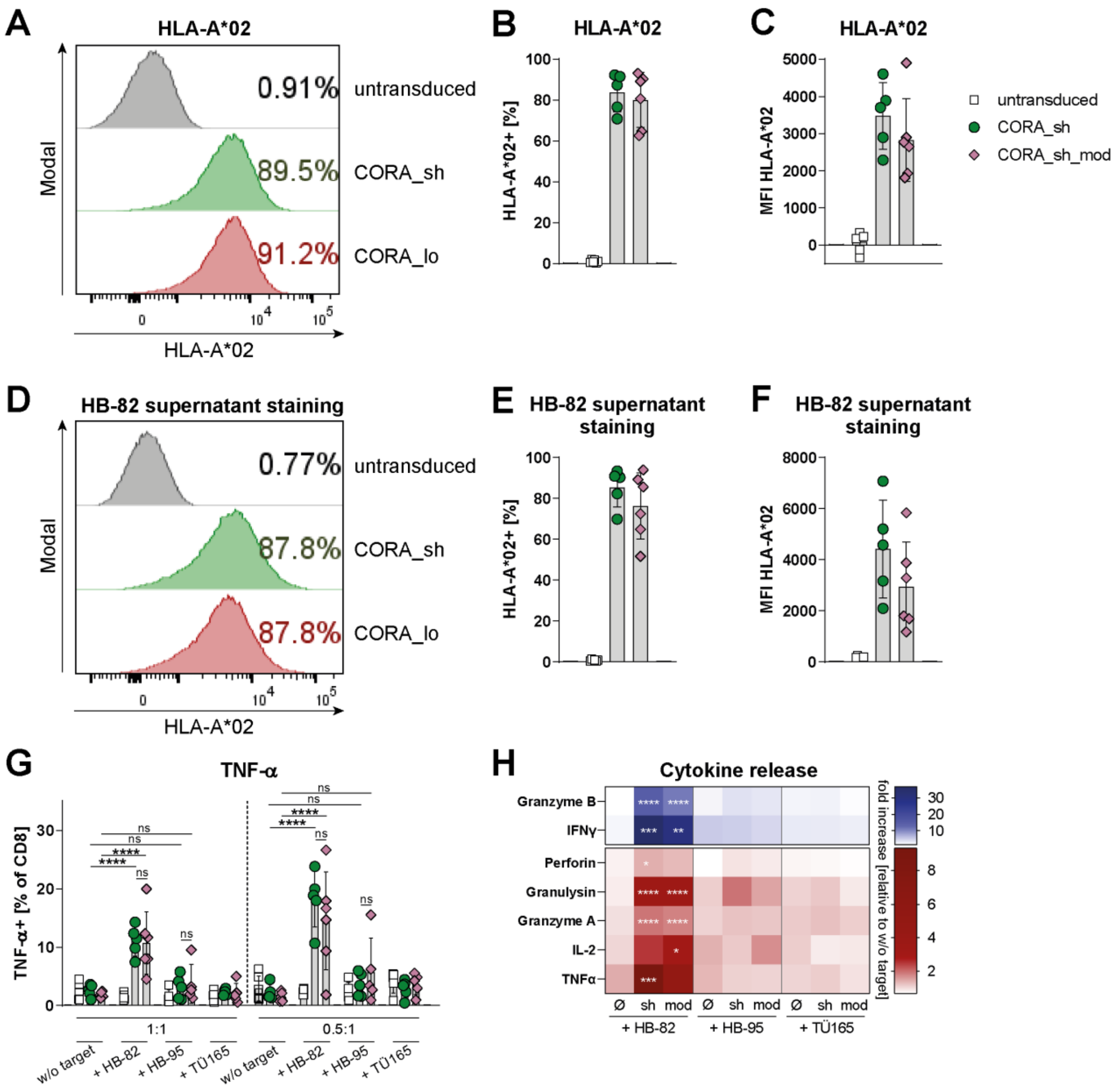


**Fig. S5: CORA-Ts with a truncated HLA-A*02 molecule as recognition domain exhibit effective and target-specific cytokine expression, which is unaltered by modification for abrogation of T-cell-sensitization.** CORA_sh receptors comprising either a truncated wildtype or a modified (CORA_sh_mod) HLA-A*02 molecule were transduced into primary CD8$^{+}$ T cells isolated from healthy donors. Untransduced T cells served as controls. **(A-F)** The HLA-A*02 component of the CORA receptor was detected by flow cytometry and staining with **(A-C)** an anti-HLA-A*02 antibody or **(D-F)** cell culture supernatant of HB-82 cells. Expression is shown as **(A, D)** representative histograms, **(B, E)** frequency (n=5-6) or (C, F) MFI (n=5-6) of CD8$^{+}$ T cells. **(G, H)** Generated CORA-Ts were cultured without (w/o) target cells or with the indicated target cells in **(G)** the indicated effector-to-target (E:T) ratio or **(H)** an E:T ratio of 5:1 for 48 h. **(G)** Expression of intracellular cytokines in CD8$^{+}$ T cells was evaluated by flow cytometry. Data are shown as scattered dot plot with mean±SD, whereby each symbol represents an independent donor (n=5-6). Statistical analysis was performed by using Two-Way ANOVA with Tukey's multiple comparisons test. ns: not significant, *p≤0.05, **p≤0.01, ****p≤0.0001. **(H)** Release of cytokines and cytotoxic mediators by untransduced (ø) and transduced CD8$^{+}$ T cells into the supernatant was assessed by LEGENDplex. Fold increase to respective T cells cultures w/o target is shown as mean (n=7-8).

### Supplementary Methods

#### Flow cytometry

Antibodies used for flow cytometry are listed in table S2. The anti-EGFRt antibody was purchased (Erbitux; ImClone Systems) and coupled to biotin (Thermo Fisher Scientific) for further use. Staining of HLA was performed by addition of human Fc block (Beckton Dickinson) for 10 min, followed by addition of respective anti-HLA antibodies.

**Table S1.** Antibodies used for flow cytometry. Phycoerythrin (PE), Peridinin-chlorophyll-protein (PerCP), fluorescein isothiocyanate (FITC), Alexa Fluor® (AF), allophycocyanin (APC), Brilliant Violet™ (BV).

| Specificity | Antibody Clone | Fluorophore | Supplier |
|---|---|---|---|
| EGFRt | - | (biotin) | ImClone Systems |
| Streptavidin | - | PE | Thermo Fisher Scientific |
| HLA-A*02 | BB7.2 | PerCP/Cy5.5 | Biolegend |
| HLA class I | W6/32 | FITC | Bio-Rad |
| $\beta_2$-microglobulin | 2M2 | PE | Biolegend |
| mouse IgG (H+L) | polyclonal $F(ab')_2$ fragment | PE | Jackson Immunoresearch |
| CD3 | SK7, UCHT1, HIT3a | AF700, PerCP | BioLegend |
| CD8 | SK1 | BV510, AF700, APC | BioLegend |
| CD25 | BC96 | APC-Cy7, PE-Cy7, BV421 | BioLegend |
| CD69 | FN50 | BV605 | BioLegend |
| CD137 | 4B4-1 | PE-Cy7, APC | BioLegend |
| Granzyme B | GB11 | Pacific Blue | BioLegend |
| TNF-a | MAb11 | APC | BioLegend |
